# Understanding the Effect of Left Prefrontal Stimulation on Positive Symptoms of Schizophrenia: A Dynamic Causal Modeling Study of Ultra-high field (7-Tesla) Resting-state fMRI

**DOI:** 10.1101/2020.02.07.939470

**Authors:** Roberto Limongi, Michael Mackinley, Kara Dempster, Ali R. Khan, Joseph S. Gati, Lena Palaniyappan

**Author notes:** ***Correspondence***, Correspondence concerning this article should be addressed to Roberto Limongi, Robarts Research Institute, 1151 Richmond St. N, UWO, London, Ontario, Canada, N6A 5B7.

## Abstract

Repetitive transcranial magnetic stimulation (rTMS), when applied to left dorsolateral prefrontal cortex (LDLPFC), reduces negative symptoms of schizophrenia, but has no effect on positive symptoms. In a small number of cases, it appears to worsen the severity of positive symptoms. It has been hypothesized that high frequency rTMS of the LDLPFC might increase the dopaminergic neurotransmission by driving the activity of the left striatum in the basal ganglia (LSTR)—increasing striatal dopaminergic activity. This hypothesis relies on the assumption that either the frontal-striatal connection or the intrinsic frontal and/or striatal connections covary with the severity of positive symptoms. The current work aimed to evaluate this assumption by studying the association between positive and negative symptoms severity and the effective connectivity within the frontal and striatal network using dynamic causal modeling (DCM) of ultra-high field (7 Tesla) resting state fMRI in a sample of 19 first episode psychosis (FEP) subjects. We found that of all core symptoms of schizophrenia, only delusions are strongly associated with the fronto striatal circuitry. Stronger intrinsic inhibitory tone of LDLPFC and LSTR, as well as a pronounced backward inhibition of the LDLPFC on the LSTR related to the severity of delusions. We interpret that an increase in striatal dopaminergic tone that underlies delusional symptoms, is likely associated with increased prefrontal inhibitory tone, strengthening the frontostriatal ‘brake’. Furthermore, based on our model, we propose that lessening of positive symptoms could be achieved by means of continuous theta-burst or low frequency (1Hz) rTMS of the prefrontal area.

## Introduction

Neuromodulatory interventions such as repetitive transcranial magnetic stimulation (rTMS) are appropriate when antipsychotic treatment fails to control psychotic symptoms [1]. While rTMS applied to left dorsolateral prefrontal cortex reduces the burden of negative symptoms [2] in schizophrenia, a recent review has highlighted that, in some cases, high frequency left prefrontal rTMS could worsen the severity of positive symptoms [3]. In this work, we explore the mechanistic basis of this phenomenon, and expound a circuit-based model to treat delusions, which are currently not clinical targets for rTMS therapy.

When treating psychosis, the effect of rTMS strongly varies with the site of stimulation. Auditory hallucinations decrease upon high frequency stimulation (HF-rTMS) of the left temporo-parietal junction [4], while left prefrontal HF-rTMS appears effective to treat negative symptoms. In contrast, while HF-rTMS of TPJ has no effect on negative symptoms, stimulating the left DLPFC appears to worsen positive symptoms in some patients [3]. Though individual studies reporting worsening of positive symptoms [5] have not identified if this effect is specific to certain positive symptoms, the earliest anecdotes indicated a specific but brief detrimental effect on delusions [6,7]. More recent trials indicate that some positive symptoms such as excitement may indeed improve with HF-rTMS of left DLPFC [8]. Importantly, out of 11 controlled trials investigating the effect of prefrontal HF-rTMS, worsening of positive symptoms have been reported in only some, as highlighted by Kennedy et al. [3] and Marzouk et al. [9]. Similarly, the extensive literature on HF-rTMS to left DLPFC in depression does not indicate any increase in the risk of positive psychotic symptoms [10]. Taken together, these observations indicate that a subset of patients with schizophrenia is likely to have a brief exacerbation of certain positive symptoms when excitatory TMS is applied to left DLPFC.

Seminal combined TMS/positron emission tomography (PET) studies have indicated the possibility of striatal dopamine release in response to prefrontal stimulation in humans [11,12]. Carlsson [13] outlined the possibility of a pyramidal glutamatergic frontostriatal feedback accelerator circuit that facilitates striatal dopaminergic output, which, when excessive, is balanced by a GABAergic inhibitory brake circuit. Based on these studies, and in keeping with the longstanding dopaminergic hypothesis of positive symptoms of schizophrenia [14], Kennedy et al. [3] hypothesized that the apparent worsening of positive symptoms on left DLPFC HF-rTMS is likely due to disrupted prefrontal excitation-inhibition balance [15], leading to left dorsal striatal dopaminergic excess via the frontostriatal network.

In this study, we further parse the hypothesis proposed by Kennedy et al. [3] and test if the intrinsic frontal connections (reflecting the inhibition/excitation balance) and the frontal-striatal connectivity (reflecting interregional influences) relate to the severity of positive symptoms. We pursued this evaluation by using dynamic causal modeling (DCM) [16] of ultra-high field (7 Tesla) resting state fMRI in a minimally medicated, actively symptomatic sample of patients with first episode psychosis (FEP). As briefly summarized below, DCM emerges as a physiologically realistic tool for evaluating this assumption, thus explaining the mechanistic basis of symptom exacerbation after prefrontal HF-rTMS and on the other hand providing insight into the parameters of left DLPFC stimulation that can be therapeutically beneficial for the same symptoms.

### Dynamic Causal Modeling of Network Connectivity

In dynamic causal modelling, differential equation models of “neural states” are fit to timeseries data. In the case of fMRI, the main goal of DCM is to make inferences about the neural causes of the BOLD signals in a-priori defined brain regions observed during either resting state [17] or task execution [16]. This allows the researcher or clinician to quantitatively infer how the timeseries are generated by (unobserved) neural activity of coupled neuronal populations.

At present, DCM is considering the most physiologically grounded technique to infer the effective connectivity between brain regions [18]. Importantly, model parameters are estimated via Bayesian inference [19]. Therefore, researchers can fit different models (each model representing one hypothesis about how the fMRI data were generated) [20,21] and crucially how the effective connectivity covaries with, for example, symptoms severity and clinical interventions (e.g., rTMS).

In this work, we capitalized on the utility of DCM to answer why positive symptoms of schizophrenia could exacerbate upon applying HF-rTMS to the left DLPFC. We fit eight models corresponding to eight core symptoms of PANSS-8 scale to resting-state fMRI data obtained at ultra-high (7T) field from 19 first episode psychosis (FEP) subjects. Using Bayesian model selection, we determined which model (i.e., symptoms metric) better explains the between-subjects variability in connections. Given the anecdotal evidence as discussed above, we expected the severity of positive symptoms, rather than negative symptoms, to relate to the frontostriatal effective connectivity [22]. In addition, given that only a subset of patients is likely to experience exacerbated positive symptoms, we aim to identify the connectivity pattern that can predict this adverse outcome of prefrontal HF-rTMS in schizophrenia.

## Methods and Materials

### Subjects

Nineteen FEP subjects participated in the study (Table 1). This patient sample has been previously reported in [23]. Subjects were recruited from the “Prevention and Early Intervention Program for Psychosis” in London, Ontario. Criteria for inclusion in the FEP group included (i) first clinical presentation with psychotic symptoms and (ii) DSM-5 [24] criteria A for schizophrenia satisfied (ii) less than 2 weeks of lifetime antipsychotic exposure. Roughly 40% of patients were not exposed to any antipsychotic at the time of assessment (Table 2). Defined Daily Dose exposure was calculated for these patients and the mean antipsychotic Defined Daily Dose in our sample was 1.05 (suggesting the average patient had had 1-day worth of the minimum maintenance dose at the time of assessment).

**Table 1.** Demographic information of participants. (C, Caucasian), (A / ME, Asian or Middle Eastern), (NSSEC, national statistics socioeconomic status), (SES, socioeconomic status), (SOFAS, social and occupational functioning assessment scale), (N/A non reliable information). Age FEP male M = 21.75, SD = 3.59; age FEP female M = 21.71, SD = 3.15.

| Subject | Ethnicity | SOFAS | Cannabis Use (self-endorsed) | Parental SES (NSSEC) | Age at Study Date | Gender | Age Psychosis Onset (years) | Duration of Untreated Psychosis (months) |
| --- | --- | --- | --- | --- | --- | --- | --- | --- |
| 1 | C | 40 | 1 | 4 | 19 | Male | 17 | 24 |
| 2 | C | 37 | 1 | 5 | 20 | Male | 20 | 4 |
| 3 | C | 40 | 1 | 2 | 19 | Male | 19 | 1 |
| 4 | C | 60 | 1 | 2 | 17 | Male | 17 | 2 |
| 5 | C | 30 | 1 | 4 | 18 | Male | 18 | 2 |
| 6 | C | 51 | 0 | 4 | 17 | Female | 16 | 9 |
| 7 | A / ME | 34 | 0 | 5 | 24 | Male | 23 | 12 |
| 8 | C | 50 | 1 | 2 | 21 | Male | 21 | 1 |
| 9 | C | 25 | 1 | 2 | 25 | Male | 20 | 59 |
| 10 | A / ME | 40 | 1 | 2 | 28 | Male | 28 | 3 |
| 11 | C | 33 | 1 | 2 | 20 | Female | 18 | 14 |
| 12 | C | 65 | 0 | 3 | 23 | Female | N/A | N/A |
| 13 | A / ME | 25 | 1 | 2 | 23 | Male | 22 | 6 |
| 14 | C | 44 | 1 | 3 | 24 | Female | 24 | 1 |
| 15 | C | 20 | 0 | 2 | 23 | Female | N/A | N/A |
| 16 | C | 55 | 1 | 4 | 20 | Male | 19 | 9 |
| 17 | C | 50 | 1 | 1 | 27 | Male | 21 | 72 |
| 18 | C | 40 | 0 | 5 | 26 | Female | 26 | 1 |
| 19 | C | 45 | 1 | 4 | 19 | Female | 19 | 1 |

**Table 2.** Participants medication at the day of assessment. (N/A non reliable information)

| Subject | Days of Antipsychotic | Antipsychotic | Dose (mg) | Duration (days) | Defined Daily Dose (DDD) |
| --- | --- | --- | --- | --- | --- |
| 1 | 0 | 0 | 0 | 0 | 0 |
| 2 | 0 | 0 | 0 | 0 | 0 |
| 3 | 7 | Paliperidone | 6 | 7 | 7 |
| 4 | 0 | 0 | 0 | 0 | 0 |
| 5 | 7 | Aripiprazole | 5 | 7 | 2.6 |
| 6 | 0 | 0 | 0 | 0 | 0 |
| 7 | 0 | 0 | 0 | 0 | 0 |
| 8 | 0 | 0 | 0 | 0 | 0 |
| 9 | 7 | Paliperidone | 3, 6 | 3, 4 | 5.5 |
| 10 | 0 | Olanzapine | 10 | 1 | 1 |
| 11 | 0 | 0 | 0 | 0 | 0 |
| 12 | 5 | Risperidone | 1.5 | 5 | 1.5 |
| 13 | 0 | 0 | 0 | 0 | 0 |
| 14 | 7 | Olanzapine | 7.5 | 7 | 5.25 |
| 15 | 0 | 0 | 0 | 0 | 0 |
| 16 | 0 | 0 | 0 | 0 | 0 |
| 17 | 0 | 0 | 0 | 0 | 0 |
| 18 | 7 | Risperidone | 1 | 7 | 1.4 |
| 19 | 10 | Aripiprazole | 5 | 10 | 3.333 |

All patients with FEP received a consensus diagnosis from 3 psychiatrists (LP/KD and the primary treatment provider) after approximately 6 months on the basis of the best estimate procedure, as described in [25], and the Structured Clinical Interview for DSM-5. All patients satisfied criteria for Schizophrenia-Spectrum Disorders, with 15 patients satisfying DSM-5 criteria for Schizophrenia and 3 for Schizoaffective Disorder. One subject lacked follow-up clinical data at 6 months, with the available baseline data suggesting a diagnosis of Schizophreniform Disorder. We use the term FEP to describe the patient group to capture all the schizophrenia-spectrum disorders as above. Informed consent from participants was obtained according to the approval by Western University’s Human Ethics Committee. Symptoms assessment was performed using Positive and Negative Syndrome Scale-8 items version [26, Table 3].

**Table 3.**
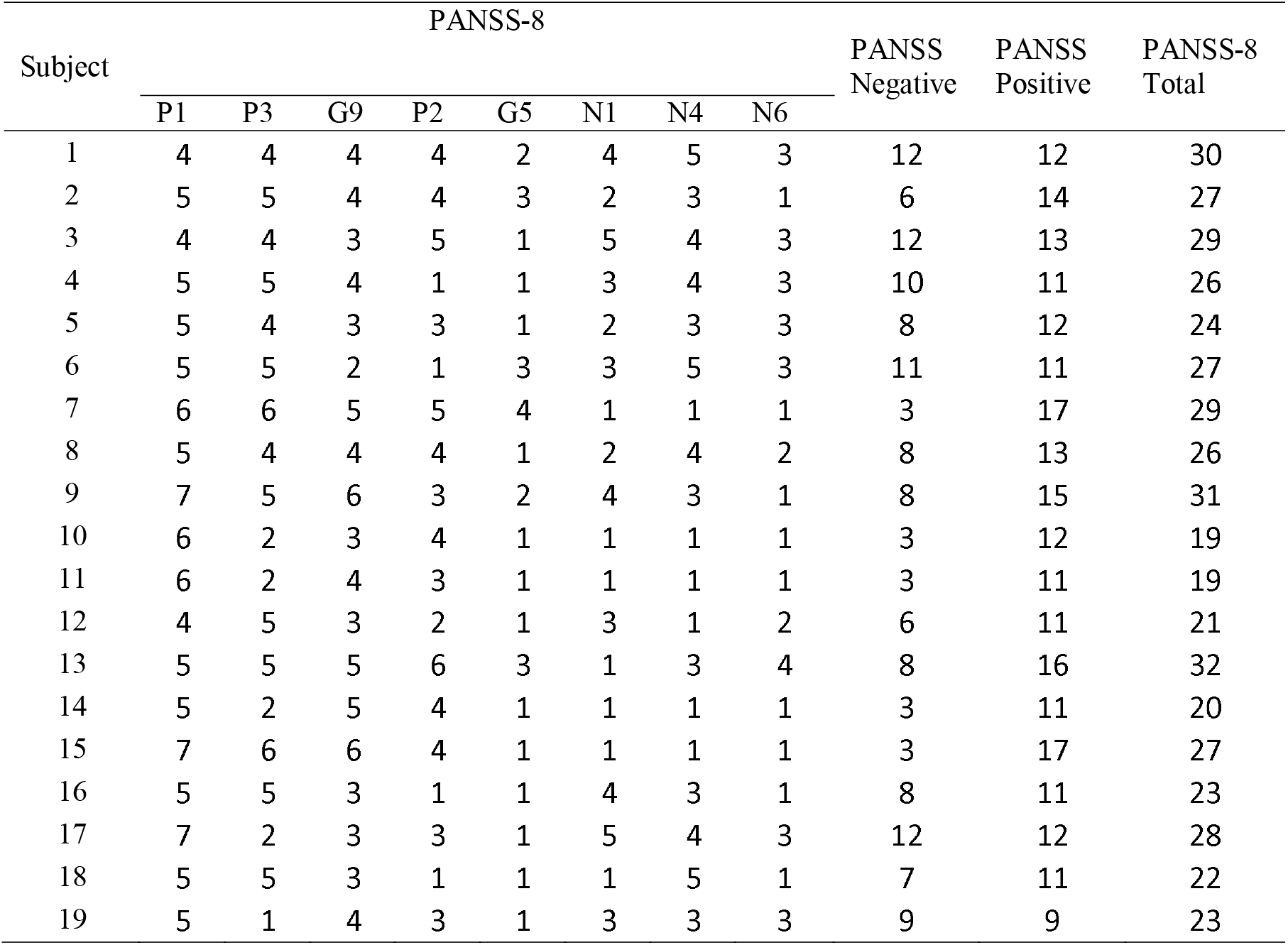
Symptoms scores assessed via the 8-item positive and negative symptoms scale (PANSS-8). HC subjects scored 1 in all metrics (P1, delusions; P2, conceptual disorganization; P3, hallucinations; N1, blunted affect; N4, social withdrawal; N6, lack of spontaneity; G5, mannerisms; G9, unusual thoughts).

### Resting-state fMRI

All data was acquired using a 680 mm neuro-optimized 7 T MRI (Siemens MAGNETOM Plus, Erlangen, Germany) equipped with an AC84 II head gradient coil and an 8-channel Tx, 32-channel Rx radiofrequency coil. We acquired 360 resting-state whole-brain functional images over 6 minutes. A gradient echo-planar-imaging sequence was used with an echo time (TE) = 20 ms, repetition time (TR) = 1000 ms, flip angle = 30 deg, field of view = 208 mm, voxel dimension = 2 mm isotropic and 63 contiguous slices. The EPI data were accelerated using GRAPPA = 3 and a multi-band factor = 3. A 3D T1-weighted MP2RAGE anatomical volume (TE/TR = 2.83/6000 ms, TI1/TI2 = 800/2700 ms) at 750 μm isotropic resolution were acquired as an anatomical reference.

### Effective Connectivity

We estimated the resting-state effective connectivity within the LDLPFC-LSTR network by fitting a two-state spectral DCM [27] to the fMRI data [17]. Functional images were realigned, normalized to the Montreal Neurological Institute (MNI) space, and spatially smoothed using a 4 mm (full width at half maximum) Gaussian kernel. We fit a general linear model to the images and included six head movement parameters and time series corresponding to the white matter and cerebrospinal fluid as regressors. We included a cosine basis set with frequencies ranging from 0.0078 to 0.1 Hz [28]. Images were high-pass filtered to remove slow frequency drifts (< 0.0078 Hz).

We identified regions with blood oxygen level fluctuations within frequencies ranging from 0.0078 to 0.1 Hz [28] via an F-contrast. We extracted the timeseries that summarized the activity within spheres (8-mm radius) in the LDPFC (MNI coordinates x= −35.5 y= 49.4 z = 32.4 [29] and in the LSTR (MNI coordinates *x*□=□-13, *y*□=□15, and *z*□=□9), [30,31]. Two-state DCM assumes excitatory and inhibitory populations of neurons within a region. Each population comprises self-inhibition connections (which are fixed parameters). Crucially, two free parameters are fit to the fMRI data: interregional excitatory-to-excitatory connections and within-region inhibitory-to-excitatory connections (Fig. 1).

For each participant, a fully connected DCM, with no exogenous inputs (c.f., task fMRI) was specified and inverted using spectral DCM. At a group level, we estimated parameters of eight parametric empirical Bayes (PEB) models [32,21] aiming to answer which of the 8 core symptoms measured using PANSS-8 scale best explained the effective connectivity within the two-node network. Models were labeled as follows: hallucinations, delusions, blunted affect, social withdrawal, disorganization, mannerism, unusual thoughts, and lack of spontaneity. We relied on Bayesian model selection procedures to select the winning model.

**Figure 1.**
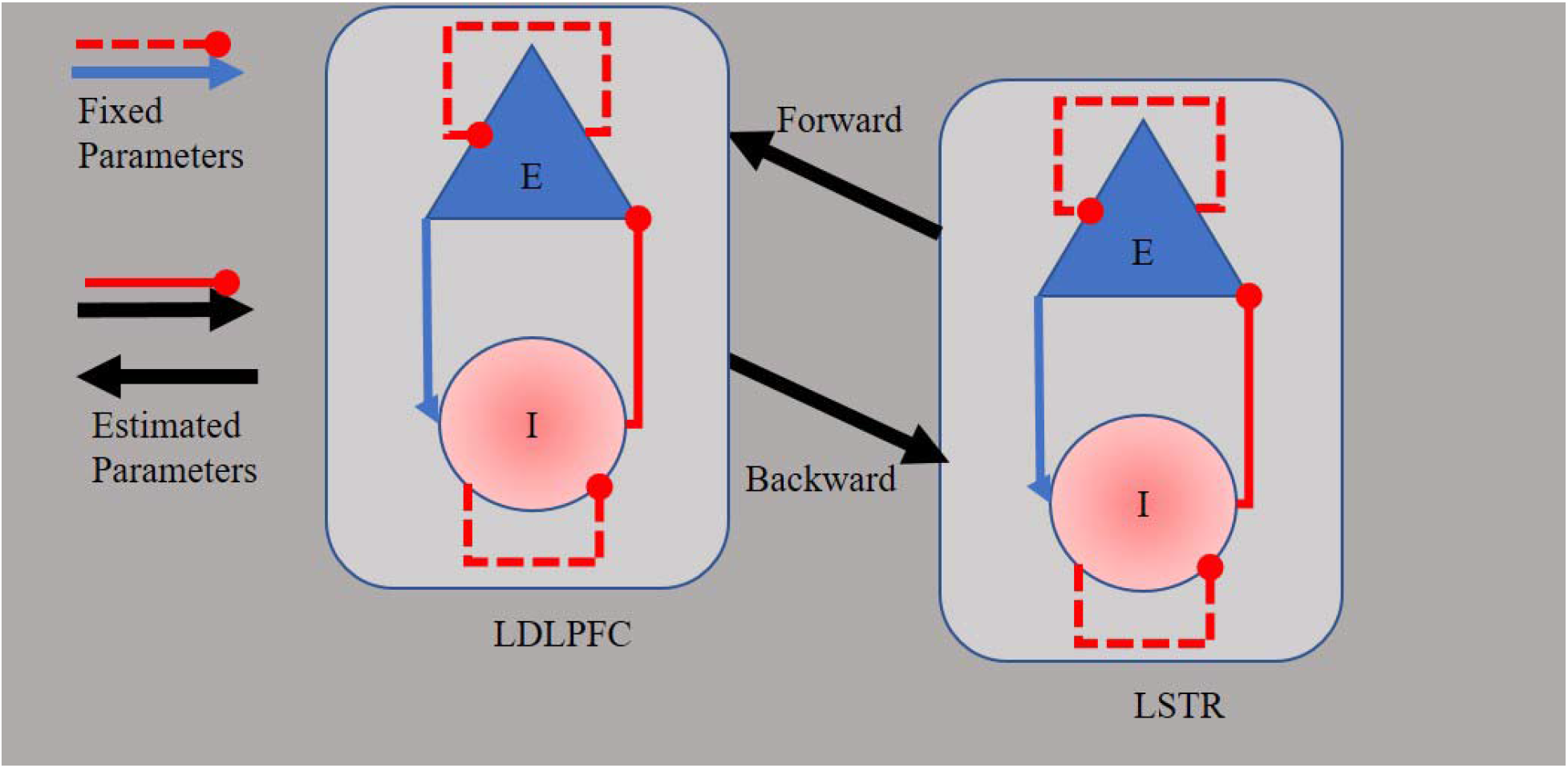
Two state dynamic causal model of the fronto striatal network. Each region comprises one population of excitatory neurons (E) and one population of inhibitory neurons (I). Parameters of effective connectivity represent the influence of inhibitory to excitatory connections (IE, assumed to be GABAergic neurons), the influence of excitatory to inhibitory connections (EI), the influence of self-inhibitory connections within each population (SE, SI), and the influence of excitatory population of one region on the excitatory population of the other region (EE, assumed to be glutamatergic connections). Whereas EI, SE, and SI parameters are fixed in the model, IE and EE are free to vary, evaluated parameters.

## Results

Bayesian model comparison revealed that the model representing the effect of delusions on the network’s effective connectivity outperformed all other models PP = 1 (Figure 2). Table 4 shows the model diagnostics in terms of the percentage of explained variance, M = 25.01 (SD =14.22). A value of at least 10 % explained variance (averaged across subjects) is considered acceptable [20]. We confirmed this via a one sample t test aiming to reject a null value of 10 %; t(_18_) = 4.6, p < .001.

**Table 4.** PEB Model diagnostics

| Subject | % of Variance<br>Accounted for by the<br>Model |
| --- | --- |
| 1 | 38.08 |
| 2 | 13.44 |
| 3 | 44.32 |
| 4 | 25.54 |
| 5 | 13.14 |
| 6 | 7.22 |
| 7 | 10.25 |
| 8 | 1.78 |
| 9 | 35.28 |
| 10 | 50.95 |
| 11 | 25.25 |
| 12 | 43.86 |
| 13 | 20.08 |
| 14 | 37.83 |
| 15 | 30.3 |
| 16 | 27.59 |
| 17 | 11.42 |
| 18 | 11.02 |
| 19 | 27.87 |

**Figure 2.**
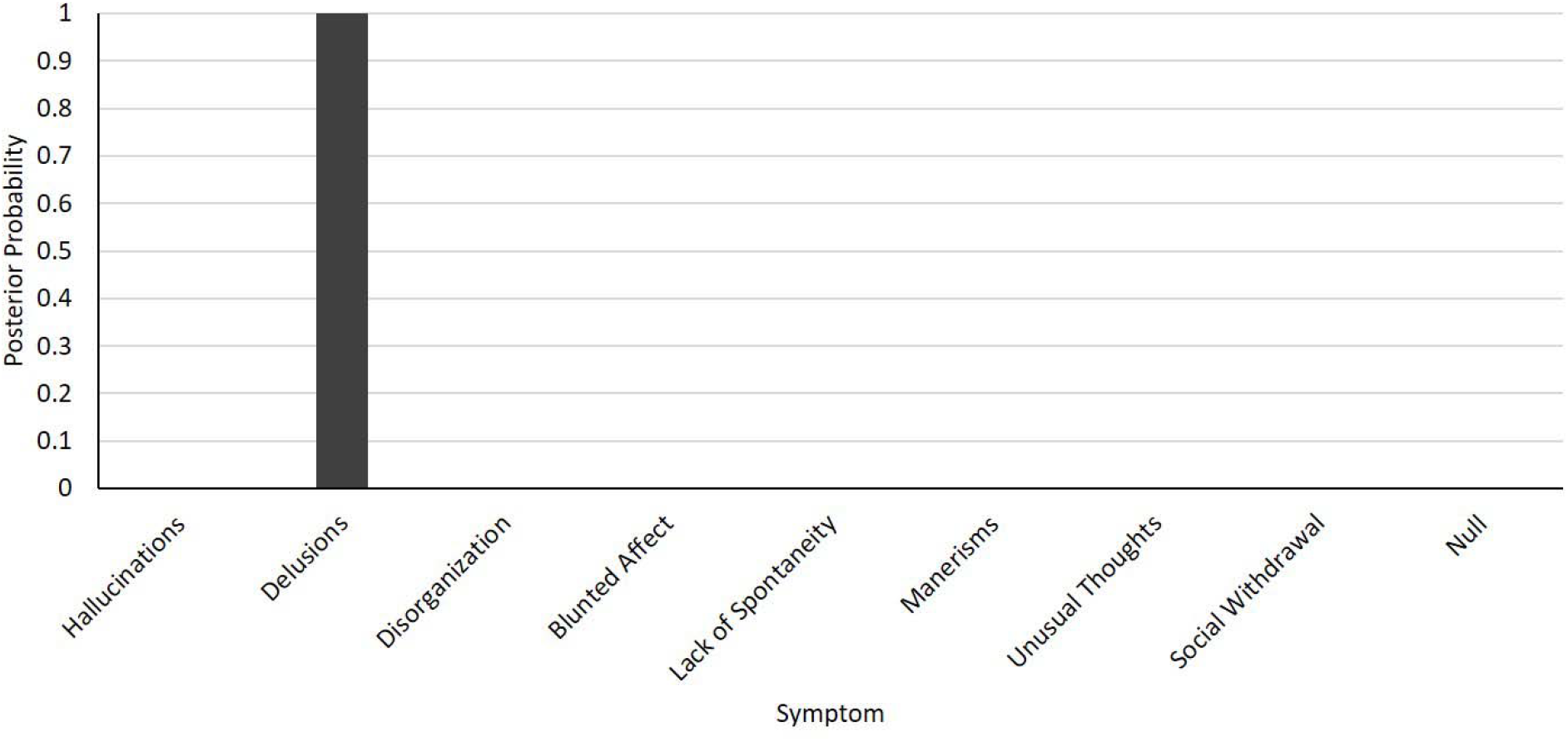
Model space and Bayesian model comparison results. Each model represented the association between one type of symptoms and the effective connectivity within the fronto striatal network. The null model represented no association between symptoms and connections.

At a group level, the model revealed positive activity of inhibitory connections (i.e. inhibition of regional excitatory neuronal pool) in both the LDPFC and the LSTR regardless of symptoms severity. Table 5 shows the parameter estimates at a subject level, and Figure 3 shows group averages (PP > 0.95). Interestingly, parameter estimates of bidirectional exogenous connections were negative, revealing bidirectional negative influence. Crucially, the LDPFC→LSTR connectivity strength decreased with the severity of delusions.

**Table 5.** Parameter estimates of connectivity strength at a subject level

| Subject | Connection |  |  |  |
| --- | --- | --- | --- | --- |
|  | LDLPFC→LDLPFC | LSTR→LDLPFC | LDLPFC→LSTR | LSTR→LSTR |
| 1 | 1.897 | -0.155 | -0.165 | 2.047 |
| 2 | 0.927 | -0.167 | -0.263 | 1.988 |
| 3 | 1.933 | -0.154 | -0.161 | 2.015 |
| 4 | 1.298 | -0.185 | -0.271 | 2.043 |
| 5 | 0.909 | -0.162 | -0.260 | 2.133 |
| 6 | 0.641 | -0.140 | -0.248 | 2.130 |
| 7 | 0.776 | -0.156 | -0.270 | 2.164 |
| 8 | 0.249 | -0.097 | -0.204 | 2.109 |
| 9 | 1.931 | -0.156 | -0.153 | 1.913 |
| 10 | 2.015 | -0.156 | -0.156 | 2.027 |
| 11 | 1.340 | -0.185 | -0.259 | 1.993 |
| 12 | 2.010 | -0.163 | -0.158 | 1.923 |
| 13 | 1.132 | -0.169 | -0.267 | 2.025 |
| 14 | 1.955 | -0.157 | -0.153 | 1.934 |
| 15 | 2.017 | -0.176 | -0.160 | 1.827 |
| 16 | 1.729 | -0.159 | -0.184 | 2.044 |
| 17 | 0.986 | -0.201 | -0.279 | 1.855 |
| 18 | 0.796 | -0.199 | -0.332 | 2.082 |
| 19 | 1.725 | -0.166 | -0.196 | 2.052 |

**Figure 3.**
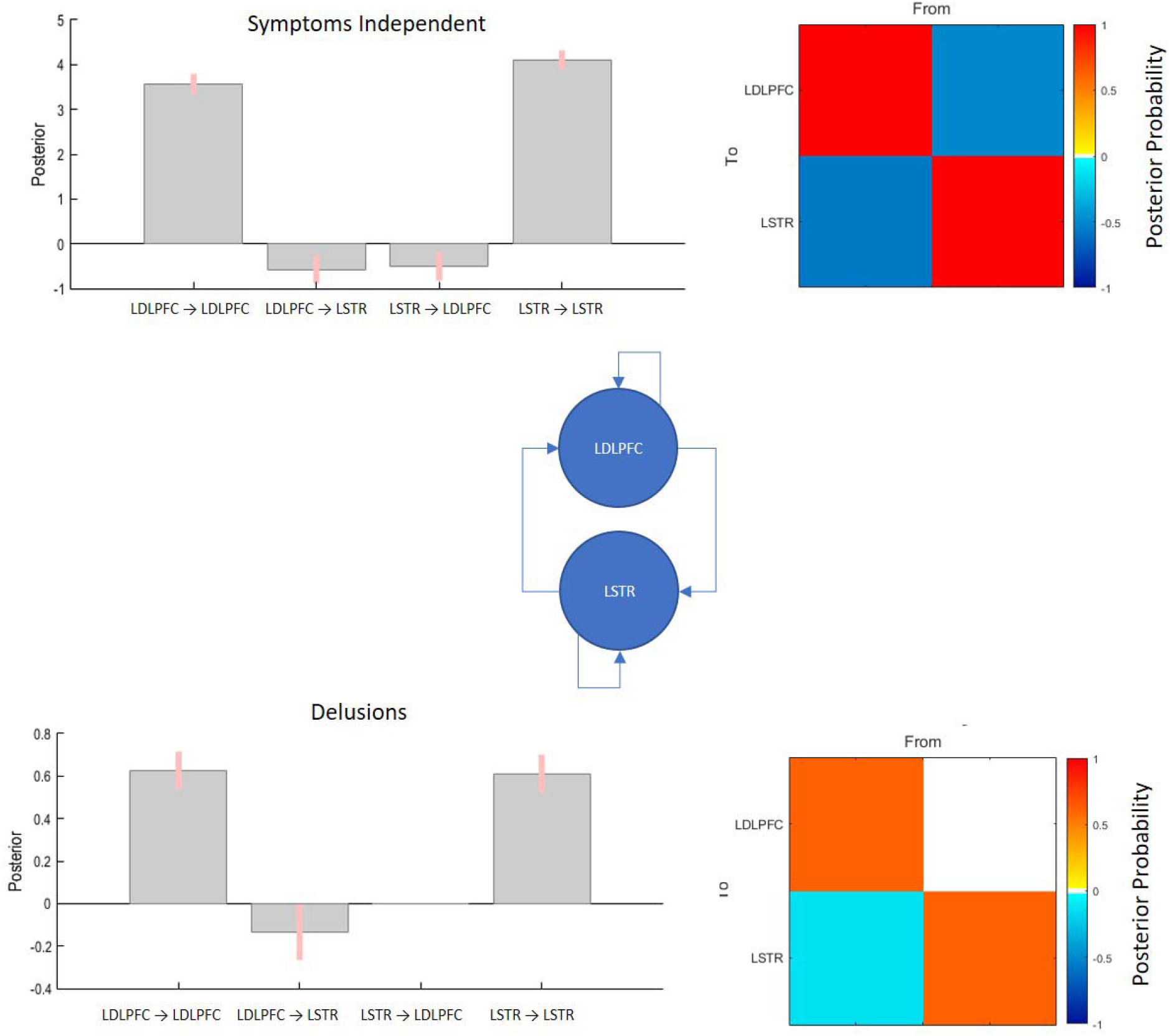
Parameter estimates of the winning (delusions) model. Top bar graph represents the mean values of parameter estimates regardless of the effect of delusions. Bottom bar graph represents the rate of change of connectivity strength as a function of severity of delusions. Left matrices represent the posterior probabilities of the parameter estimates.

## Discussion

The current work aimed to evaluate the assumption that the frontal-striatal connectivity underlies the severity of positive symptoms of schizophrenia, thus providing an empirical framework to design rTMS paradigms that can effectively reduce positive symptoms. We have 3 major findings; (1) In the absence of task-related demands (i.e. at ‘rest’), the forward and backward exogenous connections between LDLPFC and LSTR have a net inhibitory effect in patients with schizophrenia (2) Of the various characteristic symptoms measured using PANSS-8, delusions were the feature associated with the observed frontostriatal connectivity model (3) Patients with more severe burden of delusions had a strong inhibitory tone within the LDLPFC and LSTR and pronounced negative influence of the LDLPFC on the LSTR. As we elaborate upon below, these observations clarify the likely nature of glutamate-dopamine interactions in frontostriatal circuit in schizophrenia and provide a framework for future TMS studies to address treatment-resistant delusions.

We observed a bidirectional, predominantly negative influence in the exogenous connections between LDLPFC and LSTR in our sample. This indicates that at resting state, the indirect GABA-mediated frontostriatal ‘brake’ pathway is likely to be dominant [33,34], with any increase in prefrontal excitation poised to reduce striatal activity. This observation is supported by combined PET/MRS studies indicating an inverse relationship between prefrontal glutamate and striatal dopamine levels in healthy controls [35] as well as patients with schizophrenia [36]. This physiological state of frontostriatal inhibition also suggests that when dopaminergic excess occurs in the striatum, enhancing prefrontal ‘brakes’ on the striatum may have a desirable effect of reducing striatal hyperdopaminergia. As a corollary, any reduction in frontostriatal inhibition may have the effect of switching from ‘brake’ to ‘accelerator’ mode, further enhancing the hyperdopaminergic state [37].

In patients with severe delusions, we noted an increase in prefrontal and striatal inhibitory tone, with more pronounced frontostriatal inhibition (Figure 4). This pattern is consistent with enhanced, GABA-mediated, indirect frontostriatal pathways (i.e., stronger brakes). Given that this sample is of unmedicated first episode subjects who responded well to antipsychotics in the next 6 months (16 out of 19 achieved >50% reduction in overall PANSS-8 scores when clinically followed-up), we favor the inference that the observed connectivity patterns are secondary to higher striatal dopaminergic activity. Thus, enhanced self-inhibitory tones of LDLPFC and LSTR, and a strong frontostriatal ‘brake’ are likely to be compensatory changes, albeit inefficient. In this case, reducing LDLPFC’s inhibitory tone (i.e. disinhibition via NMDA blockade), may release the brakes and likely worsen delusions. This framework provides a mechanistic explanation as to why those rTMS protocols that cause disinhibition (e.g. >5-10Hz) may worsen delusions. While this conjecture is based on the glutamate hypothesis [38–41,23], our data neither supports nor refutes the primacy of glutamate over dopaminergic dysfunction in psychosis.

**Figure 4.**
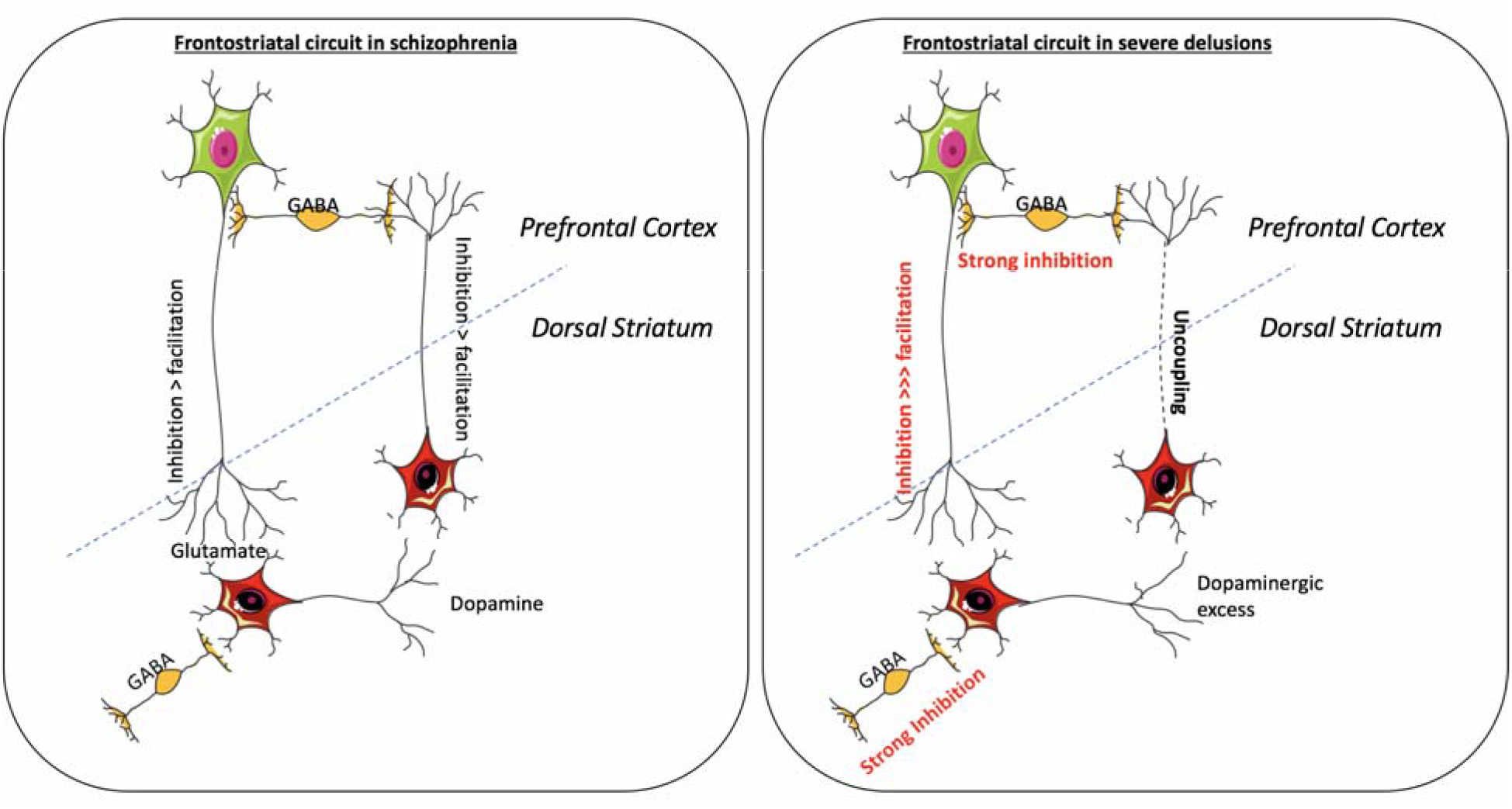
Frontostriatal connectivity and delusions. Panel A represents the bidirectional inhibitory exogenous connections presumably of glutamatergic (from dorsolateral prefrontal cortex) and dopaminergic (from striatum) nature, and GABAergic inhibitory neuronal populations within both structures. Panel B represents the state of the circuit in response to striatal hyperdopaminergic tone, with stronger inhibitory tone in both striatal and prefrontal inhibitory neuronal populations. Furthermore, the frontostriatal brakes are now stronger, and this may be essential to achieve a satisfactory response to treatment of delusions. Illustrations developed from the images provided by https://smart.servier.com/.

### Inhibitory rTMS in the LDLPF could lessen delusions symptoms

The DCM results lead us to propose that protocols that increase the inhibitory tone on cortical pyramidal neurons, may reduce further striatal DA output and thus reduce delusions. We have previously demonstrated that intermittent theta-burst stimulation (iTBS) reduced GABA/glutamate ratio in LDLPFC in healthy controls, tilting the balance towards reduced self-inhibitory tone [42]. Thus, LDLPFC iTBS can potentially increase overall excitatory output from the stimulated cortex thus abolishing the “brakes”. In contrast, continuous TBS has been shown to increase GABA (albeit in the motor cortex) [43]. Provided cTBS has the same effect on LDLPFC GABA levels in patients, this could reduce the severity of delusions in schizophrenia. Another possible protocol is increasing inhibitory tone via low frequency rTMS (e.g., 1Hz) [44], Whether the inhibitory effect of cTBS and 1Hz rTMS translates to stronger frontostriatal inhibitory control is an empirical question that could be addressed via DCM in future studies.

### DCM could provide insights on treatment efficacy

Previous works combining rTMS and DCM have shown the feasibility of DCM in describing (non-invasively) the effect of induced brain responses on the Parkinsonian brain [28] and stroke patients [45]. Other connectivity methods could also be used to estimate changes in connectivity parameters of resting state fMRI after stimulation [46]. For example, using functional (i.e., correlational) connectivity analysis Vercammen et al. [47] showed an increase in the correlation between the activity of the LTPJ and the right insula after six days of low frequency (1 Hz) rTMS. However, unlike functional connectivity analysis, DCM would provide realistic biological interpretation of connectivity parameters by explaining the direct (i.e., causal) effects of rTMS on both (within region) inhibitory (i.e., GABAergic) connections and (between region) excitatory (glutamatergic) connections. PEB models of the sort used in this work could also serve as a tool to investigate the variability of the intervention associated with demographic and antipsychotic medication. Finally, at the subject level, DCM could provide timely information to the clinician about the effectiveness of the intervention before symptoms assessment (which is time consuming).

### Limitations

The interpretation of the current results relies on several simplifying assumptions. We focused our analysis on a two-node network which does not represent the actual underlying mechanism of psychosis. For example, the severity of negative symptoms may relate to the LDPFC’s connectivity with a different seed, not evaluated here. Furthermore, the two state DCM approach comprises only two neuronal populations, which does not represent the actual canonical microcircuit either in the cortex or in the striatum. We focused on a target that is immediately accessible for rTMS and did not study the extended circuitry including the hippocampal excitatory connections that may have a critical “accelerator” effect on the striatum. Individual variability to non-invasive brain stimulation is still poorly understood; our work raises the question of utilizing frontostriatal connectivity parameters to estimate the physiological state of the corticofugal striatal pathways, before choosing the parameters of brain stimulation in schizophrenia. We expect that such an approach will pave way for expanding TMS treatment to hitherto understudied conditions such as delusional disorders and psychotic depression.

## ACKNOWLEDGMENTS

We thank Dr. Tushar Das, Mr. Trevor Szekeres, Mr. Peter Jeon and Dr. Jean Theberge for their assistance in data acquisition and archiving. We thank Dr. William Pavlovsky for consultations on clinical radiological queries. We thank Drs. Raj Harricharan, Julie Richard, Priya Subramanian and Hooman Ganjavi and all staff members of the PEPP London team for their assistance in patient recruitment and supporting clinical care. We gratefully acknowledge the participants and their family members for their contributions. Requests for data should be addressed to Dr. Lena Palaniyappan

